# *N*-acetylchitohexaose induces immune priming against *Pseudomonas aeruginosa* infection in the silkworm, *Bombyx mori*

**DOI:** 10.1101/2025.07.15.664963

**Authors:** Kazuhiro Mikami, Hiroto Nakajima, Masaki Ishii, Daisuke Yamanaka, Fumiaki Tabuchi, Masashi Muroi, Koichi Makimura, Shinya Ohata, Atsushi Miyashita

## Abstract

Lysine Motif (LysM) domain–containing receptors are evolutionarily conserved pattern recognition receptors (PRRs) that serve as key mediators of glycan sensing and innate immune activation in plants and mammals. In invertebrates, however, their role in activating innate immunity remains poorly understood, although some evidence for immunosuppressive functions exists. In this study, we performed *in silico* structural analyses and identified a putative *B. mori* LYSMD3 homolog (XP_004933441.1). This protein exhibits high structural similarity in the LysM domain to human LYSMD3, with a root-mean-square deviation (RMSD) of 0.559 Å, indicating close structural alignment. RNA-seq analysis of hemocytes isolated from silkworm larvae injected with *N*-acetylchitohexaose (GN6), a chitin-derived oligosaccharide and known ligand of human LYSMD3, revealed transcriptional activation of innate immune effectors, including antimicrobial peptide (AMP) genes such as *cecropins*. GN6 also induced *cecropin* transcription in isolated hemocytes *in vitro*, and western blotting of hemolymph confirmed elevated Cecropin B protein levels. Furthermore, GN6 and chitin significantly improved survival against *P. aeruginosa* infection, with median effective doses (ED₅₀) values of 0.62 and 0.48 µg/larva, respectively. In contrast, *N*-acetylglucosamine (GlcNAc) and shorter oligosaccharides (GN2–GN5) were ineffective. These findings provide the first molecular-level evidence of a putative glycan receptor in silkworms based on structural similarity to known LysM domains. Moreover, GN6-induced antimicrobial peptide expression and enhanced infection resistance demonstrate immune priming in this model, supporting an evolutionarily conserved glycan-sensing pathway in invertebrates.

## INTRODUCTION

The development of new therapeutic modalities independent of antibiotics and vaccines has become an urgent global priority due to the growing threat of antimicrobial resistance and the limited efficacy of existing vaccines (1). In this context, strategies aimed at enhancing innate immunity to strengthen host resistance represent universally applicable approaches that may transcend species barriers. Invertebrates have long been considered to lack adaptive immunity (2, 3). However, recent studies have reported a phenomenon termed *immune priming*, in which prior exposure to a pathogen enhances resistance to subsequent infections with the same pathogen (4). Immune priming has been observed in a variety of invertebrate species, including silkworms, fruit flies, mosquitoes, wax moths, woodlice, shrimp and mollusks, and is associated with the induction of resistance against a broad range of pathogens, including bacteria, fungi, and viruses (5–16). In some cases, this response exhibits pathogen-specific protection and immune memory-like effects that persist through metamorphosis, highlighting *immune priming* as a distinctive immunological concept that extends beyond the conventional framework of innate immunity (10, 17, 18).

We have previously demonstrated that pre-injection of heat-killed microbial cells or peptidoglycan in silkworms (*Bombyx mori*) induces the expression of the antimicrobial peptide (AMP) Cecropin B and enhances resistance against infections caused by enterohemorrhagic *Escherichia coli* and *Pseudomonas aeruginosa* (9). Furthermore, administration of heat-killed *P. aeruginosa* or *Candida albicans* confers resistance to *P. aeruginosa* but not to *C. albicans*, indicating that silkworms exhibit pathogen-specific enhancement of resistance, which may reflect underlying immune priming mechanisms (10).

Lysine Motif (LysM) domain–containing receptors are widely conserved pattern recognition receptors (PRRs) involved in glycan recognition across kingdoms, suggesting a fundamental role in innate immune systems throughout evolution (19). In plants, transmembrane LysM-type receptors recognize chitin and *N*-acetylchitohexaose (GN6), a hexamer composed of *N*-acetylglucosamine (GlcNAc) residues derived thereby triggering immune responses against fungal and bacterial pathogens (20–22). Similarly, in mammals, GN6 induces innate immune responses and enhances resistance to infections by pathogens such as *Pseudomonas aeruginosa* and *Listeria monocytogenes* (23, 24). Recent studies identified the mammalian transmembrane receptor LYSMD3 as a key sensor of GN6 and related chitin oligosaccharides. This receptor activates NF-κB signaling independent of classical TLR2 and TLR4 pathways and leads to robust cytokine production (e.g., IL-6 and IL-8) (19).

These observations support the hypothesis that LysM-mediated glycan recognition is functionally conserved in both plant and animal immune systems. Within invertebrates, the role of LysM domain-containing receptors in immune activation remains poorly understood and appears paradoxical. The transmembrane receptor immune active (*ima*), a *Drosophila* homolog of mammalian *LYSMD3/4*, recognizes peptidoglycan and modulates the Imd–Relish pathway (25). Unlike the immune-stimulatory LysM receptors in plants and mammals, *ima* primarily functions as a negative regulator to prevent excessive immune activation. Thus, it remains unknown whether insect possess LysM domain–containing receptors capable of immune activation, similar to those in plants and mammals. Clarifying this gap could reveal evolutionarily conserved immune pathways extending from plants and mammals to insects, underscoring the broader significance of glycan sensing mechanisms.

To explore this possibility further, we focused on the *B. mori*, an experimentally tractable insect model widely used in innate immunity studies (26). However, whether *B. mori* possesses structurally conserved LysM domain–containing receptors with similar functional roles remains unknown. In this study, we first performed *in silico* structural analyses to determine whether *B. mori* possesses a LYSMD3-type receptor homologous to the mammalian counterpart. We identified a candidate receptor that shares a conserved LysM domain architecture with human LYSMD3. Given this structural similarity and the established role of LYSMD3 in GN6 recognition and innate immunity activation in mammals (19), we hypothesized that GN6 might similarly stimulate immune responses in insects through a conserved LysM-mediated pathway. To test this, we evaluated whether GN6 could induce antimicrobial peptide expression and enhance infection resistance in *B. mori*, aiming to uncover a conserved mechanism of glycan-mediated immune activation across species.

## RESULTS

### 1. Identification and structural analysis of a LYSMD3 homolog in *B. mori*

To investigate whether *Bombyx mori* possesses a homolog of mammalian LYSMD3, we searched the NCBI protein database for annotated LysM domain–containing proteins. Human LYSMD3 (NP_938014.1) is annotated as “LysM and putative peptidoglycan-binding domain-containing protein 3,” and a *B. mori* (XP_004933441.1) bearing the same annotation pattern was identified as a putative homolog. We then compared the amino acid sequences and predicted structures of human LYSMD3 and the silkworm protein. Sequence alignment revealed conservation of the LysM domain in both proteins (Figure 1A), suggesting a shared chitin-binding module. Structural alignment of the predicted LysM domain regions further demonstrated a high degree of similarity, with a root-mean-square deviation (RMSD) of 0.559 Å (Figure 1B), indicating that local domain architecture is well conserved between the two species.

**Figure 1.**
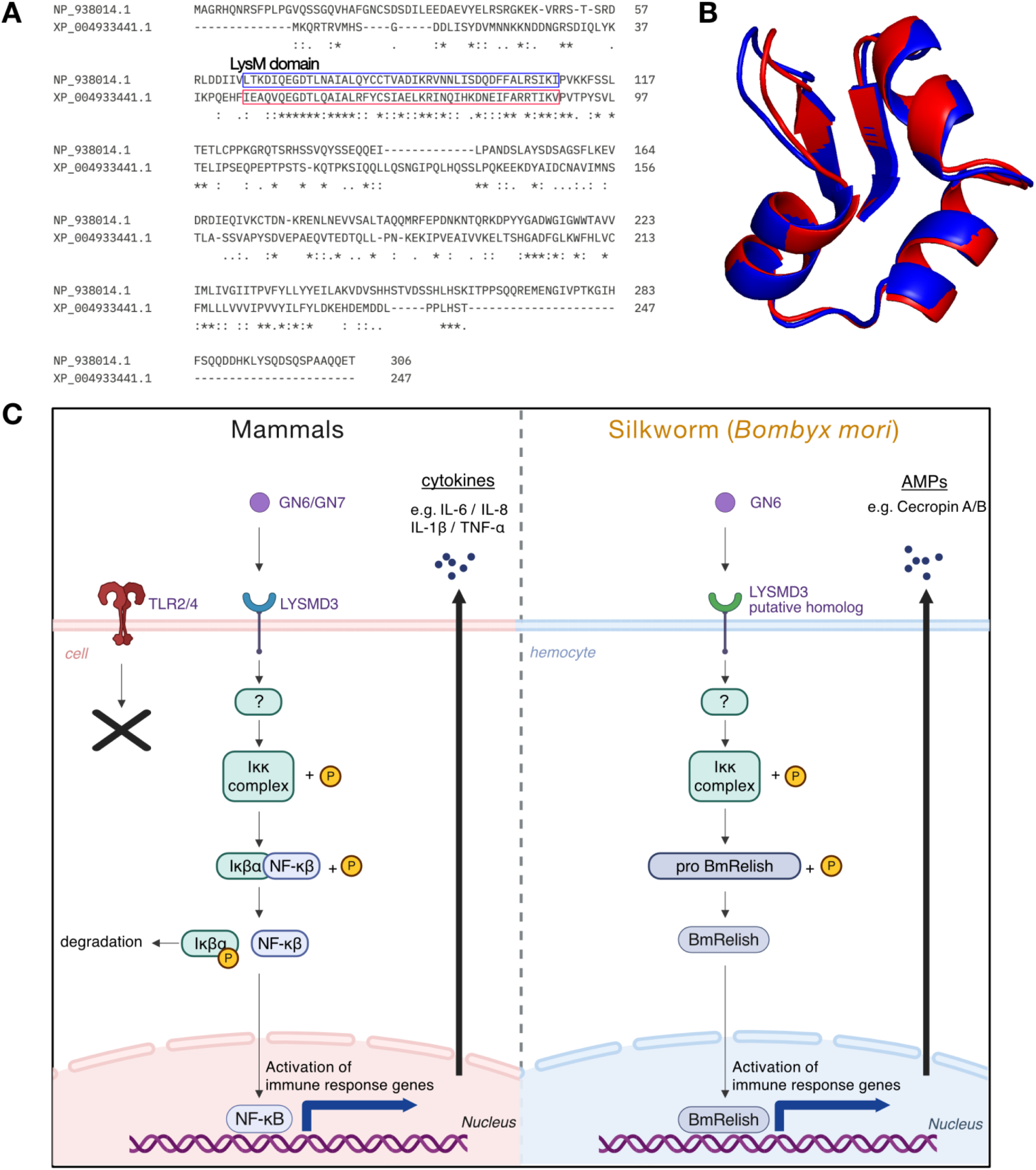
Structural comparison and conserved signaling model of LysMD3 proteins in humans and *Bombyx mori*. (A) Multiple sequence alignment of human LYSMD3 (NP_938014.1) and the *B. mori* homolog (XP_004933441.1) was performed using CLUSTAL Omega (version 1.2.4). The LysM domain in the human sequence (residues 65–109) and the predicted LysM domain in the *B. mori* homolog (residues 45–89) are indicated by blue and red boxes, respectively. Asterisks (*) denote fully conserved residues, colons (:) indicate strong similarity, and periods (.) represent weaker similarity in amino acid properties. (B) Predicted 3D structures corresponding to the LysM domain regions were obtained from the AlphaFold Protein Structure Database and visualized using PyMOL after membrane configuration assignment via CHARMM-GUI. The human model is shown in blue and the *B. mori* model in red. Structural superimposition revealed a high degree of similarity, with an RMSD of 0.559 Å. (C) Hypothetical model of GN6-mediated innate immune activation via LysM-type receptors in mammalian and insect systems. Left: In mammalian cells, GN6 is recognized by LYSMD3, which activates the IκB kinase (Ikk) complex, leading to phosphorylation and degradation of IκBα. This allows NF-κB to translocate into the nucleus and induce immune-related gene expression. Right: In *B. mori* hemocytes, the LYSMD3 homolog is presumed to recognize GN6 and activate the Ikk complex, which in turn activates the *Bombyx* Relish homolog (BmRelish), inducing the expression of antimicrobial peptides such as Cecropin A and B. These pathways share a conserved feature of glycan recognition via LysM domains and suggest that GN6-mediated innate immune activation may be evolutionarily conserved. The schematic was created using BioRender.com.

In mammals, GN6 has been shown to bind LYSMD3 and induce proinflammatory cytokine production, and *N*-acetylchitoheptaose (GN7) have been demonstrated to elicit LYSMD3-dependent cytokine production in murine macrophages (19). LYSMD3 also binds fungal cell wall components more strongly than TLR2 or TLR4, suggesting a primary role in fungal recognition (19). In *Drosophila*, the LYSMD3/4 homolog *immune active* (ima) modulates the Imd–Relish pathway (25), an innate immune cascade triggered by bacterial elicitors such as peptidoglycan and mediated by the NF-κB–like factor Relish. This pathway is functionally conserved in *B. mori* as the Imd–IKKβ–Relish cascade (27). Based on these observations, we hypothesized that GN6 activates the NF-κB/Relish pathway via a LysM-type receptor in *B. mori*, representing a potentially evolutionarily conserved mechanism of innate immune activation (Figure 1C). We therefore proceeded to investigate the functional activity of GN6 in mammalian cultured cells and silkworm immune systems.

### 2. GN6 induces proinflammatory cytokine expression in RAW264.7 cells

GN6 has been previously reported to activate innate immune responses in the murine macrophage cell line RAW264.7, inducing proinflammatory mediators such as *Il1b* and *Tnf* (28). To verify that the commercially sourced GN6 used in this study exhibited similar activity, we assessed its effects in the same cell line. RAW264.7 cells were cultured and stimulated with GN6 at various concentrations for 2 hours, followed by RNA extraction and quantitative real-time PCR (qRT-PCR) analysis of cytokine gene expression. GN6 upregulated the expression of the proinflammatory cytokine genes *Il1b* and *Tnf*, in a dose-dependent manner (Supplemental Figure 2). These results supported the use of GN6 in the subsequent experiments to evaluate its immune-activating effects in the silkworm.

### 3. GN6 does not activate TLR2- or TLR4-mediated NF-κB signaling in a reporter assay

To examine whether GN6 activates NF-κB signaling via human TLR2 or TLR4, we performed a luciferase reporter assay using HEK293 cells transfected with expression plasmids for either receptor and an NF-κB–responsive luciferase construct. Cells were stimulated with GN6, and NF-κB activation was assessed by measuring luciferase activity. GN6 treatment did not induce significant NF-κB–dependent luciferase activity in either TLR2- or TLR4-expressing cells compared to unstimulated controls (Figure 2). In contrast, the positive controls lipoteichoic acid (LTA, 1 µg/mL) and lipopolysaccharide (LPS, 1 µg/mL) robustly activated NF-κB signaling through TLR2 and TLR4, respectively, confirming the responsiveness of the reporter system. These results indicate that neither TLR2 nor TLR4 alone is sufficient to mediate NF-κB activation in response to GN6 under the conditions tested.

**Figure 2.**
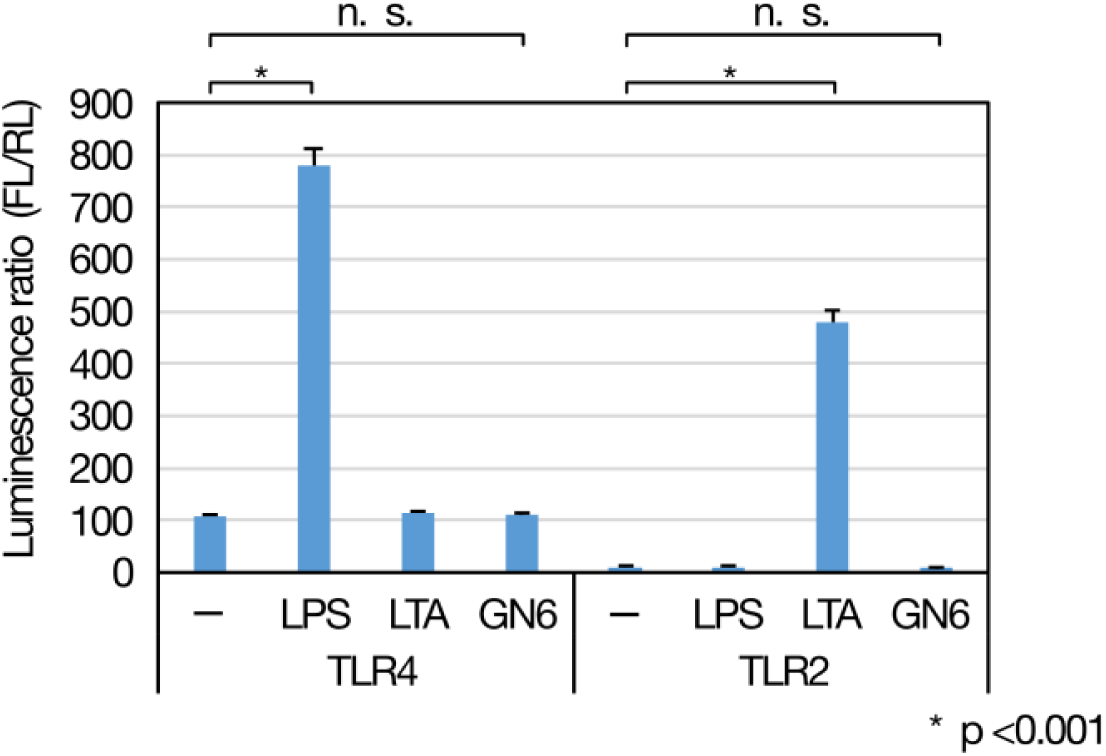
GN6 does not activate TLR2 or TLR4 signaling pathways. NF-κB activation was assessed by a luciferase reporter assay using HEK293 cells stably expressing either human TLR2 or TLR4/CD14/MD2. Cells were stimulated with GN6 (5 µg/mL) for 6 h, and firefly luciferase (FL) and Renilla luciferase (RL) activities were measured. Relative luciferase activity was calculated as the FL/RL ratio. Lipopolysaccharide (LPS, 1 µg/mL) and lipoteichoic acid (LTA, 1 µg/mL) served as positive controls for TLR4 and TLR2 activation, respectively. GN6 failed to induce NF-κB activation in either TLR2- or TLR4-expressing cells, whereas both positive controls elicited significant activation (*p < 0.001). Data represent mean ± SD (n = 3). n.s., not significant.

### 4. GN6 induces innate immune gene expression in *B. mori* hemocytes

Having established that GN6 activates mammalian macrophages independently of TLR2 or TLR4, we next investigated whether it also induces innate immune responses in *B. mori*. To address this, we performed transcriptome analysis of hemocytes isolated from saline or GN6-injected larvae, followed by differential gene expression and Gene Ontology (GO) enrichment analyses to identify immune-related transcriptional changes. Transcriptome analysis identified 59 genes significantly upregulated and one gene significantly downregulated in response to GN6 injection (adjusted p-value < 0.05, |log₂FoldChange| ≥ 1; see Supplementary Table 1). The upregulated genes included representative immune effectors such as cecropins, peptidoglycan recognition proteins, and proteases. The only downregulated gene was annotated as gamma-glutamyl transpeptidase (γ-GT). GO enrichment analysis revealed significant over-representation of immune-related terms, including antimicrobial peptide (AMP) activity and defense response (Figure 3A). These findings demonstrate that GN6 activates innate immune gene expression in silkworm hemocytes, suggesting conservation of chitin oligosaccharide recognition pathways between insects and mammals.

**Figure 3.**
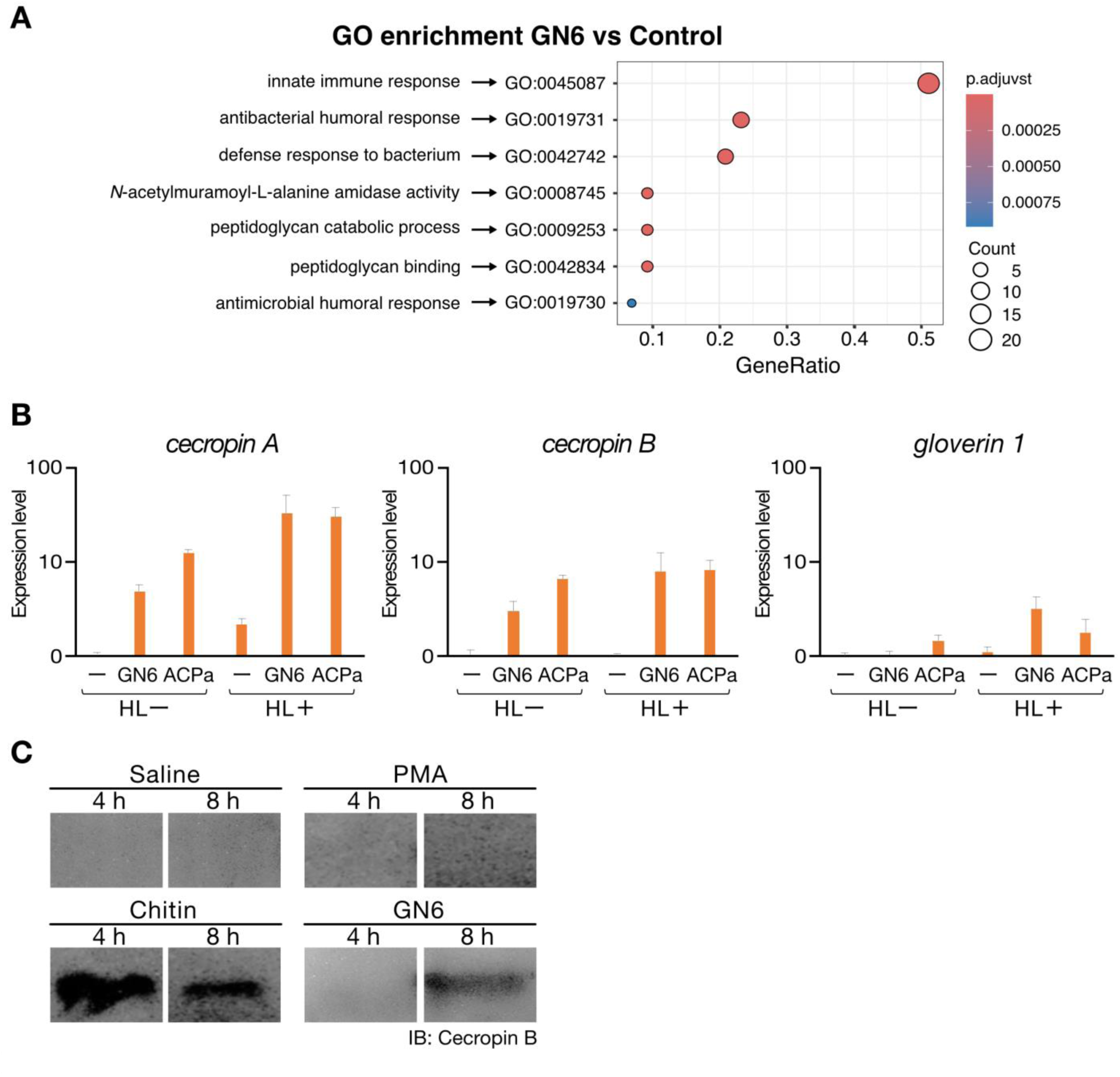
GN6-induced transcriptional and translational responses in *Bombyx mori* hemocytes and hemolymph. (A) Gene Ontology (GO) enrichment analysis of GN6-responsive differentially expressed genes (DEGs). Dot plot shows enriched GO terms among DEGs upregulated by GN6 treatment in silkworm hemocytes. Dot size indicates the number of DEGs associated with each GO term, color scale represents adjusted p-values (p.adjust), and the x-axis indicates the ratio of DEGs annotated to each term (GeneRatio). (B) GN6-induced expression of antimicrobial peptide genes in B. mori hemocytes. Hemocytes isolated from day 4 fifth-instar larvae were cultured overnight and stimulated with GN6 (0.05 mg/mL) or autoclaved Pseudomonas aeruginosa (ACPa), with or without 20% hemolymph (HL+ and HL−, respectively). After 3.5 h, total RNA was extracted, and expression levels of antimicrobial peptide genes (Cecropin A, Cecropin B, and Gloverin 1) were measured by qRT-PCR, normalized to *Eif4* expression. Results show fold induction relative to unstimulated controls. Addition of hemolymph enhanced transcriptional responses, especially for Cecropin A and Cecropin B. Data represent mean ± SEM (n = 3), shown on a logarithmic scale. (C) Induction of Cecropin B protein in *B. mori* hemolymph following GN6 or chitin treatment. Fifth-instar day 2 larvae were injected with saline, PMA (0.5 ng/larva), chitin (2.5 µg/larva), or GN6 (2.5 µg/larva) and incubated at 27°C for 4 or 8 h. Hemolymph proteins were concentrated by ethanol precipitation and analyzed by SDS-PAGE and Western blotting using anti-Cecropin B antibody. Cecropin B was specifically detected in GN6- and chitin-treated samples but absent in saline and PMA controls.

### 5. GN6 induces AMP gene expression in *B. mori* hemocytes

To investigate whether GN6 directly stimulates immune cells in *B. mori* to induce AMP gene expression, such as *cecropin A, cecropin B* and *gloverin 1*, we conducted an *ex vivo* culture assay using hemocytes isolated from fifth-instar day 4 larvae. The target genes included *cecropin A*, *cecropin B*, and *gloverin 1*, which are key effectors in the insect innate immune response. Hemocytes were preincubated overnight and stimulated the following day with either GN6 or heat-killed *Pseudomonas aeruginosa* PAO1 (ACPa). After 3.5 hours of stimulation, total RNA was extracted, and AMP gene expression was analyzed by qRT-PCR. GN6 stimulation significantly upregulated the expression of *cecropin A*, to a similar extent as ACPa treatment (Figure 3B, left). *cecropin B* expression was also significantly increased by both GN6 and ACPa treatments (Figure 3B, center). In contrast, *gloverin 1* expression was only marginally elevated by ACPa and showed no clear induction in response to GN6 (Figure 3B, right). These results suggest that GN6 activates *B. mori* hemocytes to induce the expression of specific AMPs genes, particularly members of the Cecropin family.

### 6. GN6 and chitin induce the expression of Cecropin B protein in silkworm

To assess whether GN6 or chitin induces AMP protein production *in vivo*, hemolymph was isolated from silkworms injected with each compound and analyzed for Cecropin B levels by western blotting. Protein levels were clearly elevated in GN6- and chitin-treated groups relative to the saline control group (Figure 3C). In contrast, phorbol 12-myristate 13-acetate (PMA), a known immune stimulant in mammalian systems (29), failed to induce detectable levels of Cecropin B under the same conditions. These findings indicate that both GN6 and chitin activate the innate immune system in *B. mori* at the organismal level, promoting the secretion of AMPs. The results are consistent with our previous observation of AMP gene induction in isolated hemocytes as described in the preceding section.

### 7. GN6 and chitin activate innate immune responses and induce immune priming in *B. mori*

Previous results demonstrated that GN6 and chitin upregulate AMP gene expression in silkworm hemocytes and induce Cecropin B protein production, as shown in Figure 3. To test whether GN6 injection promotes infection resistance, we employed an *in vivo Pseudomonas aeruginosa* infection model. We selected *P. aeruginosa*, a pathogen known to be susceptible to Cecropin peptides and previously shown to induce immune priming in *B. mori* following peptidoglycan pretreatment (10). Fifth-instar day 2 larvae were injected with GN6 or chitin (2.5 µg/larva) and incubated at 27°C for 4 hours, followed by infection with *P. aeruginosa* and incubation at 37°C for 18 hours. *N*-acetylglucosamine (GlcNAc), PMA, and saline were included as separate treatment groups under the same conditions. Larvae pretreated with GN6 or chitin exhibited significantly improved survival compared to the saline group (Figure 4A).

**Figure 4.**
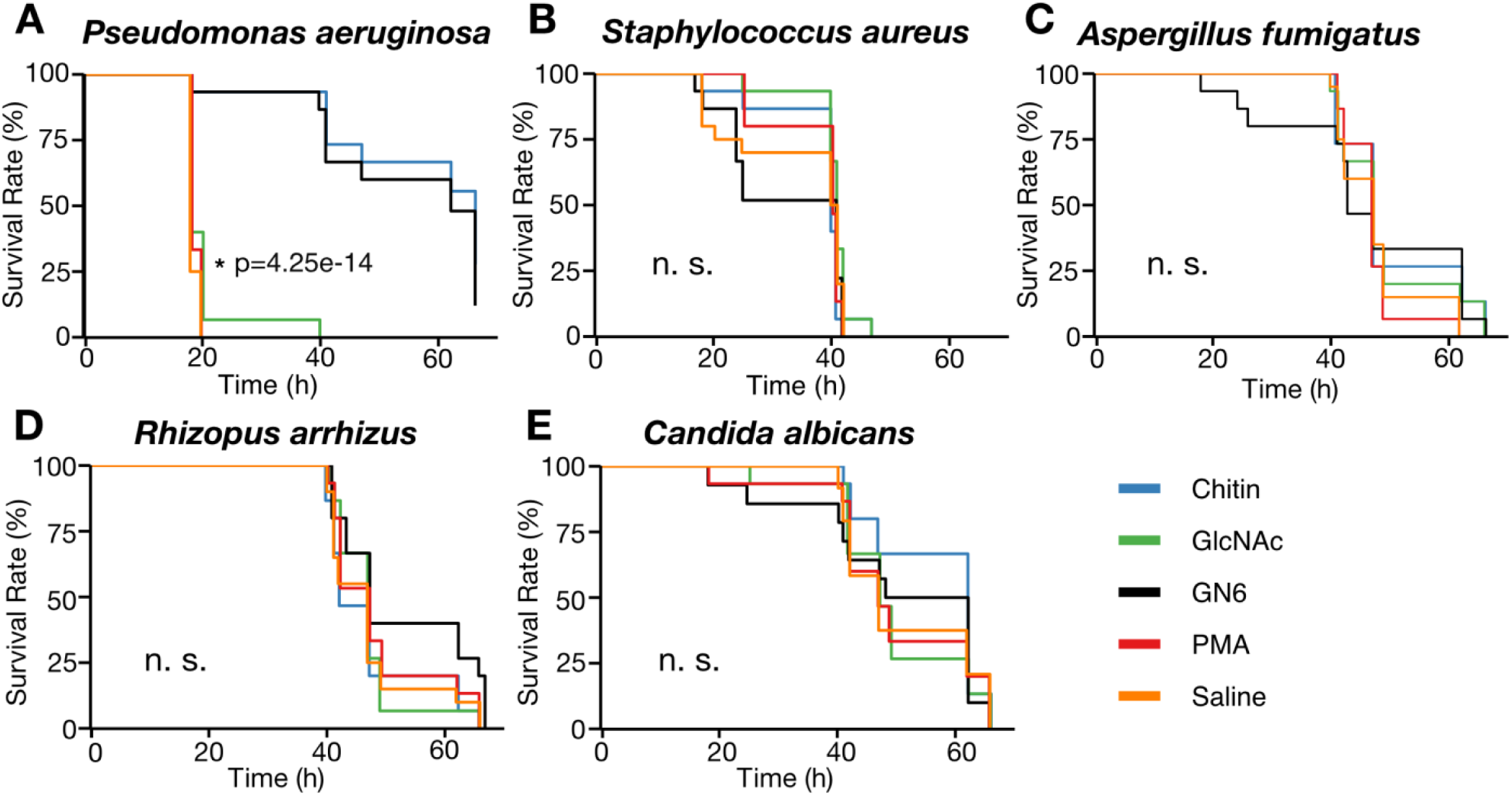
GN6 and chitin confer pathogen-specific infection resistance in *Bombyx mori*. (A) Larvae pretreated with GN6 or chitin (2.5 µg/larva) showed significantly improved survival following *Pseudomonas aeruginosa* infection compared to the saline control group (p = 4.25 × 10⁻¹⁴, log-rank test). No survival benefit was observed in larvae pretreated with GlcNAc (2.5 µg/larva) or PMA (0.5 ng/larva). (B–E) Pretreatment with GN6, chitin, GlcNAc, or PMA did not improve survival in larvae infected with *Staphylococcus aureus* (B), *Aspergillus fumigatus* (C), *Rhizopus arrhizus* (D), or *Candida albicans* (E). No statistically significant differences in survival curves were observed (n.s., p > 0.05). For all experiments, day 2 fifth-instar larvae (n = 15 per group) were injected with the test compound and challenged with the indicated pathogen 4 h later. Larvae were incubated at appropriate temperatures, and survival was monitored over time. Kaplan–Meier survival curves were generated, and group comparisons were performed using the log-rank test. Curve legend colors indicate the pretreatment condition.

### 8. GN6 and chitin enhance resistance specifically against *P. aeruginosa*, but not other tested pathogens

Having established that GN6 and chitin confer immune priming–mediated resistance to *P. aeruginosa*, we next examined whether these compounds similarly enhance resistance to other pathogens. In particular, we tested *Staphylococcus aureus*, *Aspergillus fumigatus*, *Rhizopus arrhizus*, and *Candida albicans*. *S. aureus* was included because previous silkworm studies have shown little or no immune-priming effect against this Gram-positive bacterium (30), whereas the three fungal pathogens were selected for their chitin-rich cell walls, which could differentially interact with GN6- or chitin-mediated immune pathways (31, 32). Using the same infection method and treatment conditions as with *P. aeruginosa*, we assessed larval survival following pretreatment with GN6 or chitin. In all cases, no significant improvement in survival was observed compared to the saline control (Figure 4B–E), indicating that neither compound conferred resistance to these pathogens. These results suggest that the immune priming effect of GN6 and chitin in *B. mori* is pathogen-specific, providing enhanced resistance to the Gram-negative bacterium *P. aeruginosa*, but not to the Gram-positive *S. aureus* or to the fungal pathogens tested.

### 9. Structure–activity relationship of chitin-derived oligosaccharides in infection resistance induction

To evaluate the dose-dependent effects of GN6 and chitin on infection resistance, fifth-instar day 2 larvae of silkworm were injected with GN6 or chitin at doses of 2.5, 1.25, 0.63, 0.31, or 0.15 µg/larva. Four hours after pretreatment, the larvae were challenged with *P. aeruginosa*, and survival was assessed after 18 hours. Both compounds increased survival in a dose-dependent manner (Figure 5), with calculated ED₅₀ of 0.62 µg/larva for GN6 and 0.48 µg/larva for chitin.

**Figure 5.**
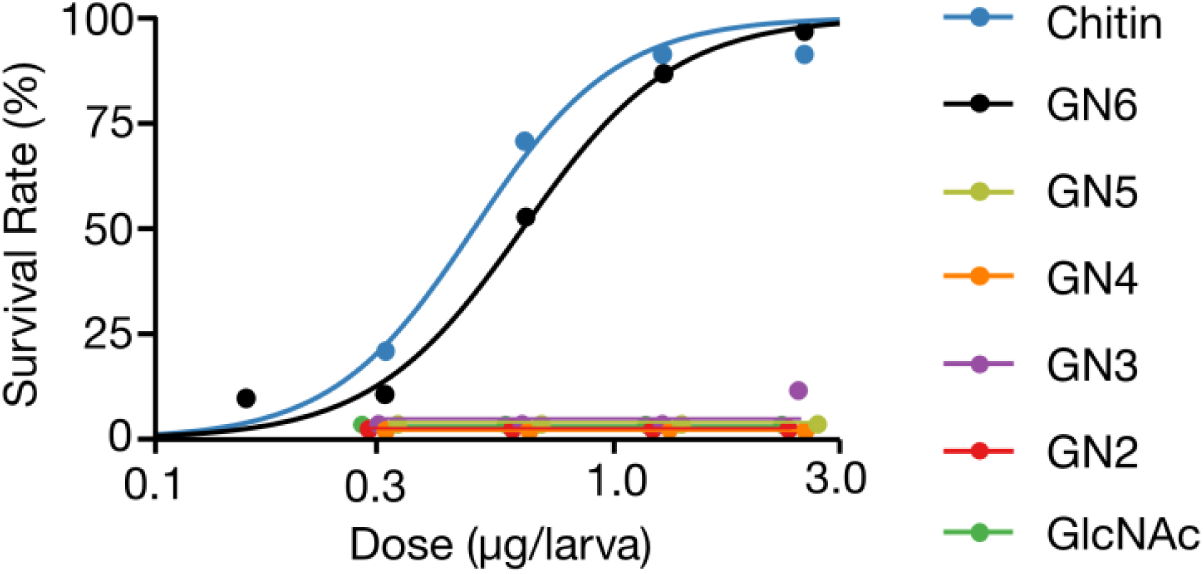
GN6 and chitin induce dose-dependent immune priming against *Pseudomonas aeruginosa*, while lower-order chitin oligosaccharides are inactive. Day 2 fifth-instar larvae were injected with GN6, chitin, or related oligosaccharides (GN2– GN5, GlcNAc) at various doses (0.15–2.5 µg/larva). After 4 h, larvae were challenged with *P. aeruginosa*, and survival was assessed 18 h post-infection (n = 15 per group). Dose– response curves were generated using nonlinear regression analysis. Both GN6 and chitin conferred infection resistance in a dose-dependent manner, with ED₅₀ values of 0.62 µg/larva and 0.48 µg/larva, respectively. In contrast, no increase in survival was observed in larvae treated with GN2–GN5 or GlcNAc, suggesting that a minimum polymer length is required to induce immune priming in *B. mori*.

We next tested whether chitin oligosaccharides with lower degrees of polymerization (GN2-GN5) and monomeric GlcNAc could induce comparable effects under the same conditions. None of these shorter oligosaccharides improved larval survival, and survival rates were comparable to those of the saline control (Figure 5). These results establish a clear structure-activity relationship, in which the induction of immune priming– associated resistance to *P. aeruginosa* in silkworms requires chitin oligosaccharides with a minimum degree of polymerization of six GlcNAc units: neither shorter oligosaccharides (GN2-GN5) nor the monomeric unit (GlcNAc) were sufficient to confer protection.

## DISCUSSION

This study provides the first evidence of the three-dimensional structural conservation of LYSMD3 in the silkworm (*Bombyx mori*), although its function remains to be characterized. We performed *in silico* structural analyses of human and silkworm LYSMD3 proteins and demonstrated a high degree of conservation in the LysM domain, the ligand-binding region. Concurrently, we showed that GN6, a known ligand of human LYSMD3 (19), induces AMP expression and immune priming against *Pseudomonas aeruginosa* infection in *B. mori*. These results suggest structural conservation of LYSMD3 and raise the possibility of a conserved mechanism for glycan-induced immune activation. Our *in silico* analysis provides a molecular basis for hypothesizing a conserved mechanism of glycan recognition across species.

GN6 stimulation of murine RAW264.7 macrophages significantly upregulated *Il1b* and *Tnf* expression, consistent with previous reports (28). In our study, GN6 did not activate NF-κB in HEK293 reporter cells expressing either TLR2 or the TLR4/CD14/MD2 complex. This observation is consistent with previous findings that recombinant TLR2 and TLR4 proteins do not bind *Alternaria alternata* spores, whereas LYSMD3 was shown to associate with fungal cell walls in a biochemical assay (19). Additionally, LYSMD3 was detected in the membrane fraction of RAW264.7 cells, and its knockdown attenuated cytokine induction upon stimulation with GN7, supporting its role as a functional glycan receptor (19). In the silkworm model, GN6 administration induced significant transcriptional upregulation of multiple genes involved in innate immunity. Gene Ontology enrichment analysis highlighted AMP activity and defense response as significantly overrepresented terms. Moreover, GN6 treatment stimulated expression of AMP genes such as *cecropins* in isolated hemocytes and increased Cecropin B protein levels *in vivo*, consistent with transcriptomic data. These immune responses likely contributed to the enhanced survival observed following *P. aeruginosa* infection, although direct causal relationships remain to be established. Among the differentially expressed genes, only γ-glutamyl transpeptidase (γ-GT) showed significant downregulation. The functional implications of this change are currently unknown. Given previous evidence that Relish, an NF-κB homolog in *B. mori*, is required for AMP gene induction (27), GN6 signaling likely activates innate immunity through the Imd–Relish pathway. Together with findings from mammalian studies, these results suggest that GN6 can activate innate immune responses in both vertebrates and invertebrates, supporting the existence of a conserved glycan-sensing pathway. However, the identity and functional role of the receptor responsible for GN6 recognition in silkworms remain to be determined. Future studies should investigate whether GN6 binds directly to the putative *B. mori* LYSMD3 homolog and whether disruption of this receptor affects downstream Relish activation and AMP production.

In this study, GN6 and chitin selectively enhanced silkworm resistance against *P. aeruginosa*, but conferred no protection against other pathogens such as *Staphylococcus aureus*, *Candida albicans*, *Aspergillus fumigatus*, or *Rhizopus arrhizus*. The lack of protection against *S. aureus* is consistent with its known resistance to cecropin-type AMPs (30). In contrast, the inability to protect against fungi, despite the presence of chitin in their cell walls, indicates that exposure to chitin oligomers alone is insufficient for initiating effective immune priming against these organisms. Although GN6 failed to confer protection against all tested fungal pathogens, it remains unclear whether this reflects a general limitation of GN6-mediated priming against fungi or if species-specific differences in fungal cell wall composition might also play a role. Further comparative analysis of fungal recognition and immune activation would be necessary to clarify this point. Identifying the molecular determinants underlying this pathogen specificity, including receptor-ligand interactions and downstream signaling specificity, represents an important objective for future research. Notably, LysM-type receptors exhibit divergent roles across species and contexts. In *Drosophila*, the *LYSMD3/4* homolog immune active (*ima*) negatively regulates the Imd–Relish pathway upon recognition of bacterial peptidoglycan, thereby suppressing excessive antimicrobial peptide induction (25). In contrast, mammalian LYSMD3 is directly binds chitin, and activates NF-κB–dependent cytokine production in response to fungal glycans (19). Given the structural similarity and the immune-activating profile observed in our silkworm model, we speculate that the identified *B. mori* LYSMD3 homolog functions more like mammalian LYSMD3 than *ima*. Nevertheless, the mechanistic basis for such divergent immune functions, despite the conservation of the LysM domain, remains unresolved. Possible explanations include differences in ligand specificity, intracellular domain structure, receptor localization, and the nature of downstream adaptor molecules. Clarifying these factors will require biochemical analyses of ligand binding, localization studies of the receptor, identification of interacting signaling components, and genetic approaches such as RNA interference or gene knockout targeting the silkworm LYSMD3 homolog.

We further investigated the structure–activity relationship of chitin-derived oligosaccharides in inducing immune responses in silkworms. GN6 and chitin significantly enhanced survival following *P. aeruginosa* infection in a dose-dependent manner, whereas shorter oligosaccharides (GN2–GN5) and monomeric GlcNAc did not confer protection. These findings indicate that a minimal structural length (six or more GlcNAc units) is required to trigger immune activation. The calculated ED₅₀ for GN6 and chitin were 0.62 µg/larva and 0.48 µg/larva, respectively, suggesting that oligosaccharides of sufficient length elicit comparable levels of immune stimulation. This size-dependent requirement is consistent with observations in mammalian systems, where chitin oligosaccharides of comparable length are needed to elicit immune responses (33).

Early studies in murine infection models (first in 1989 and later in 2003) demonstrated that chitin and its derived oligosaccharide GN6 significantly improved survival following *P. aeruginosa* infection (23, 24), although these effects were initially viewed as isolated findings due to the absence of a conceptual framework for innate immune memory. The emergence of the “*trained immunity*” and “*immune priming*” paradigms in the 2010s redefined these early results, providing a mechanistic basis for long-lasting, memory-like innate immune responses across animal phyla (34, 35). Within this revised context, our findings in the silkworm model parallel and extend the murine data, highlighting GN6 as an evolutionarily conserved immunostimulatory glycan.

Taken together, our findings furnish the first integrated structural and functional evidence that the silkworm harbors a LysM-domain receptor that is homologous to mammalian LYSMD3 and capable of sensing chitin-derived hexamers. Engagement of this receptor by GN6 triggers antimicrobial-peptide production and confers pathogen-specific protection against *P. aeruginosa*, establishing immune priming in an insect model. These results position chitin oligosaccharides and LysM-type receptors at the core of an evolutionarily ancient innate-immune module that spans invertebrates and mammals. Further investigations are needed to elucidate the molecular mechanisms underlying these observations, including direct characterization of ligand-receptor interactions and downstream signaling pathways. Collectively, this work bridges invertebrate and mammalian immunology, revealing ancient glycan-sensing pathways that may inform next-generation approaches to immune enhancement.

## MATERIALS AND METHODS

### Sequence alignment and structural prediction/comparison

The amino acid sequences of human LYSMD3 (NP_938014.1) and its homolog in *Bombyx mori* (XP_004933441.1) were retrieved from the NCBI Protein database (https://www.ncbi.nlm.nih.gov/protein/). Domain annotations were performed using InterProScan version 5.73–104.0 (https://www.ebi.ac.uk/interpro/), confirming that both proteins contain a LysM domain (ProSite profile: PS51782, Integrated: IPR018392). Multiple sequence alignment was conducted using CLUSTAL Omega version 1.2.4 (36). Based on the alignment results, the LysM domain was identified as corresponding to residues 65–109 in human LYSMD3 and residues 45–89 in the *B. mori* homolog.

Structural predictions were carried out using ColabFold v1.5.5, which implements AlphaFold2 with MMseqs2-based homology search (37). The resulting PDB files were processed through CHARMM-GUI Membrane Builder (https://www.charmm-gui.org/) to convert formats and assign membrane insertion coordinates. No molecular dynamics simulations were performed in this study; the membrane configuration was used solely to assist structural orientation and downstream visualization. Structural alignment and comparison between the human and silkworm models were performed using PyMOL version 3.1.4.1 (Schrödinger, LLC). Root mean square deviation (RMSD) values were calculated to assess structural similarity. In addition to the full-length structures, the predicted LysM domain regions identified above were extracted and analyzed separately to evaluate local structural conservation.

### Evaluation of immune-stimulatory activity of GN6 in RAW264.7 cells

To evaluate whether N-acetylchitohexaose (GN6; Cayman Chemical, Ann Arbor, MI, USA; Cat# 17864) stimulates mammalian immune cells to induce the expression of proinflammatory cytokine genes, an *in vitro* assay was performed using the murine macrophage cell line RAW264.7. Cells were seeded into 24-well plates and cultured in RPMI 1640 medium (Thermo Fisher Scientific, Waltham, MA, USA) supplemented with 10% heat-inactivated fetal bovine serum (FBS) and 1% PenStrep (Thermo Fisher Scientific) at 37°C in a humidified atmosphere with 5% CO₂ until reaching approximately 80% confluence. GN6 was then added to the culture medium at final concentrations of 2, 10, and 50 µg/mL. As a positive control, lipopolysaccharide (LPS; InvivoGen, San Diego, CA, USA) was added at final concentrations of 50, 500, and 5000 EU/mL. For the unstimulated group, only culture medium was added. After stimulation, cells were incubated for an additional 2 hours under the same conditions.

Total RNA was extracted using TRIzol reagent (Thermo Fisher Scientific), followed by isopropanol precipitation in the presence of sodium acetate and glycogen. RNA pellets were washed with 75% ethanol, air-dried, and dissolved in nuclease-free water (Qiagen, Hilden, Germany). RNA concentrations were measured using a Qubit 4 Fluorometer (Thermo Fisher Scientific). cDNA was synthesized using a commercial reverse transcription kit (ReverTra Ace, Toyobo, Osaka, Japan). Quantitative real-time PCR (qRT-PCR) was performed using Thunderbird Next SYBR qPCR Mix (Toyobo, QPX-201) on a 7500 Fast Real-Time PCR System (Applied Biosystems, Foster City, CA, USA), with MicroAmp Fast 96-well Reaction Plates (Cat# 4346907, Applied Biosystems). Primer pairs targeting *Il1b*, *Tnf*, and the reference gene *Rps18* were used. Primers were dissolved in Primer TE buffer (NIPPON GENE, Cat# 316-90025) prior to use. Gene expression levels were normalized to Rps18, and relative expression was calculated using the 2^−ΔΔCt method with the unstimulated group as the baseline (set to 1).

Each condition was tested in triplicate (N = 3). Statistical analysis was performed using one-way analysis of variance (ANOVA), followed by Tukey’s multiple comparison test. A p-value of less than 0.05 was considered statistically significant.

### TLR reporter assay using HEK293 cells

To assess whether GN6 activates human innate immune signaling via TLR2 or TLR4, a reporter assay was performed using HEK293 cells. Cells were cultured in DMEM (Thermo Fisher Scientific) supplemented with 10% fetal bovine serum and 1% PenStrep at 37°C in a humidified atmosphere with 5% CO₂. Plasmids encoding human TLR2 or TLR4/CD14/MD2, a firefly luciferase gene under the control of an NF-κB–responsive promoter, and a constitutively expressed Renilla luciferase gene were provided by a collaborator and amplified in *Escherichia coli.* Cells were transfected with the plasmid mixture using FuGENE HD transfection reagent (Promega, Madison, WI, USA) and incubated at 37°C in 5% CO₂ for 24 hours prior to stimulation.

Transfected cells were seeded into 24-well plates (Iwaki, AGC Techno Glass Co., Ltd., Japan) and stimulated with GN6 at a final concentration of 5 µg/mL. As positive controls, lipopolysaccharide (LPS, 1 µg/mL) and lipoteichoic acid (LTA, 1 µg/mL; InvivoGen, San Diego, CA, USA) were added. Phosphate-buffered saline (PBS) was used as a vehicle control for the unstimulated group.

After 6 hours of incubation at 37°C in a 5% CO₂ atmosphere, cells were lysed with Glo Lysis Buffer (Promega), and lysates were transferred to white 96-well plates. Luciferase activities were measured using the Dual-Glo® Luciferase Assay System (Promega, Cat# E2920) according to the manufacturer’s instructions. Firefly and Renilla luminescence were sequentially recorded, and firefly activity was normalized to Renilla activity (FL/RL ratio). Data are presented as fold induction relative to the unstimulated control.

Each condition was tested in triplicate (N = 3). Statistical analysis was performed using one-way ANOVA followed by Tukey’s multiple comparison test. A p-value of less than 0.05 was considered statistically significant.

### Insects

Eggs of *Bombyx mori* (silkworm; strain KINSYU × SHOWA) were obtained from Ehime-Sanshu (Ehime, Japan). The hatched larvae were reared on an antibiotic-containing artificial diet (Silkmate 2S, Nihon Nosan Corporation, Yokohama, Japan) until the fifth instar larval stage (38).

### RNA Extraction, mRNA Sequencing, and Differential Expression Analysis

Fifth-instar day-2 silkworm larvae were injected with either 2.5 µg of GN6 per larva or saline as a control. The larvae were incubated for 4 hours at 27°C following injection, after which hemolymph was collected. Hemolymph from three larvae per treatment was pooled to form one biological sample. Hemocytes were pelleted by centrifugation at 1000 × g for 5 minutes at 4°C. The supernatant was discarded, and the cell pellet was resuspended in TRIzol™ Reagent and mixed thoroughly by pipetting. The lysate was centrifuged at 13,000 rpm for 5 minutes, and the upper aqueous phase was collected. An equal volume of ethanol was added, and total RNA was extracted using the Direct-zol™ RNA MiniPrep Kit (Zymo Research, Cat# R2050), according to the manufacturer’s instructions. RNA concentration was quantified using the Qubit™ RNA HS Assay Kit, 500 assays (Thermo Fisher Scientific, Cat# Q32852).

mRNA sequencing was outsourced via Nippon Genetics Co., Ltd. to Novogene Co., Ltd., where library preparation and sequencing were performed using the Illumina NovaSeq X Plus platform, generating approximately 6 gigabases (Gb) of data per sample. Downstream analyses, including quality control, alignment, and differential gene expression analysis, were performed using an in-house *Bombyx mori* transcriptome database. Differential gene expression analysis was conducted with the DESeq2 package (version 1.43.1) in R (version 4.5.0). Differentially expressed genes (DEGs) were defined as those with an adjusted p-value (padj) < 0.05 and absolute log₂ fold change ≥ 1. Gene annotation for DEGs was assigned using SilkDB3.0 annotation files (https://silkdb.bioinfotoolkits.net), and the annotated DEG list is provided as Supplementary Table 1. Gene Ontology (GO) enrichment analysis was performed using the enricher function of the clusterProfiler package (version 4.16.0) in R. GO term annotations for *B. mori* genes were obtained from SilkDB annotation files. The statistical significance of enrichment was assessed using the Benjamini-Hochberg method to control the false discovery rate (FDR). Results were visualized using the dotplot function in clusterProfiler.

### Analysis of AMPs gene expression in cultured hemocytes from *Bombyx mori*

Hemolymph was collected from fifth-instar day 4 *Bombyx mori* larvae by cutting the abdominal prolegs. Hemocytes were washed with insect saline and suspended in IPL-41 medium supplemented with 2% fetal bovine serum and 1% PenStrep. The cell suspension was adjusted to a density of 2.7 × 10⁵ cells in 0.3 mL per well and seeded into 48-well plates (Iwaki, Cat# 3830-048). After overnight incubation at 27°C, cells were stimulated the next morning with either GN6 (final concentration: 0.05 mg/mL) or heat-killed *Pseudomonas aeruginosa* PAO1 (ACPa, 1:500 dilution). Control groups were cultured without any stimulants. For both GN6 and ACPa conditions, parallel cultures were prepared with or without the addition of 20% silkworm hemolymph to the medium. After 3.5 hours of incubation at 27°C, cells were harvested by centrifugation at 1100 × g for 7 minutes at 4°C and lysed in TRIzol reagent with vortexing.

Total RNA was extracted according to the manufacturer’s protocol. Complementary DNA was synthesized using ReverTra Ace® qPCR RT Master Mix with gDNA Remover (Toyobo, Osaka, Japan; Code FSQ-201). Quantitative real-time PCR (qRT-PCR) was performed using Thunderbird® Next SYBR® qPCR Mix (Toyobo, Osaka, Japan) on a Fast Real-Time PCR System (Applied Biosystems). Gene-specific primers targeting *Bombyx mori Eif4a*, *Cecropin A*, *Cecropin B*, and *Gloverin 1* were used. Gene expression was normalized to the housekeeping gene *Eif4a*, and relative expression was calculated by setting the expression level in the unstimulated control group to 1. No-template controls (H₂O) were included as negative controls. Each condition was tested in triplicate (N = 3). Statistical significance was assessed using one-way ANOVA followed by Tukey’s multiple comparison test, with p-values < 0.05 considered significant.

### Western blotting

The silkworm larvae were injected with PMA 10 ng/larva, chitin 2.5 µg/larva (FUJIFILM Wako Pure Chemical Corp., Osaka, Japan), GN6 2.5 µg/larva, or saline and incubated at 27°C for 4 hours. Hemolymph was collected and centrifuged to separate the supernatant. The supernatant was mixed with 99.5% ethanol at a ratio of 1:5 (v/v), followed by centrifugation. The resulting pellet was collected and dissolved in SDS sample buffer mixed with β-mercaptoethanol, then denatured by heating at 99°C for 5 minutes.

The denatured samples were subjected to SDS-PAGE using Mini-PROTEAN TGX™ 4%–20% Tris-Glycine gels (Bio-Rad Laboratories, Hercules, CA, USA) at 200 V and 40 mA. Proteins were transferred to a PVDF membrane filter at 500 V and 40 mA on ice. After transfer, membranes were blocked with 1% skim milk in buffer.

For immunodetection, membranes were blocked with 1% skim milk in Tris-buffered saline with Tween 20 (TBS-T) and incubated overnight at 4°C with a rabbit polyclonal anti-Cecropin B antibody (Abcam, Cambridge, UK; Cat# ab27571) diluted 1:1000. After washing, membranes were incubated for 3 hours at room temperature with horseradish peroxidase–conjugated goat anti-rabbit IgG (SouthernBiotech, Birmingham, AL, USA; Cat# 4030-05) diluted 1:5000 with gentle agitation. Signals were detected using Western Lightning® Plus-ECL chemiluminescent substrate (PerkinElmer, Waltham, MA, USA).

### Bacteria and fungi

For infection experiments, *Staphylococcus aureus* (MSSA1 strain) and *Pseudomonas aeruginosa* (PAO1 strain) were cultured overnight at 37°C in LB-10 medium. Full-growth cultures were diluted in sterile saline to prepare suspensions of 1×10⁶ CFU/mL for *S. aureus* and 5×10³ CFU/mL for *P. aeruginosa*. *Candida albicans* (TIMM1768 strain) was cultured in YPD medium at 37°C. Full-growth cultures were diluted in sterile saline to prepare a suspension of 1×10⁵ CFU/mL. Conidial suspensions of *Aspergillus fumigatus* (Af293 strain) and *Rhizopus arrhizus* (IFM46105 strain) were prepared by harvesting spores from potato dextrose agar and 1/10 Sabouraud dextrose agar, respectively, and adjusting the concentration to 1×10⁵ spores/mL, as described previously (39).

### Immune priming assay in *B. mori*

To evaluate the immune priming effect in *B. mori*, larvae (fifth instar, day 2) were injected with chitin, GN6, or GlcNAc at a dose of 2.5 µg per larva, each suspended in sterile saline. The control group received an equal volume of saline alone. After injection, the larvae were incubated at 27°C for 4 hours.

Following the priming treatment, larvae were infected by injection with *Pseudomonas aeruginosa* (PAO1 strain) or *Staphylococcus aureus* (MSSA1 strain), and incubated at 37°C for survival monitoring. In fungal infection experiments, larvae were similarly given infections with *Candida albicans* (TIMM1768 strain), *Aspergillus fumigatus* (Af293 strain), or *Rhizopus arrhizus* (IFM46105 strain). Larvae infected with *C. albicans* or *A. fumigatus* were incubated at 37°C, whereas those infected with *R. arrhizus* were incubated at 27°C. Survival was monitored over time, and Kaplan–Meier survival curves were generated. Statistical significance was evaluated using the log-rank test. Each experimental group was tested in triplicate (N = 3), with five silkworms per trial (n = 5 per replicate).

### Dose–response and structure–activity analysis of chitin oligosaccharides in *B. mori*

To evaluate the dose-dependent induction of infection resistance by chitin and chitin-derived oligosaccharides, *B. mori* larvae (fifth instar, day 2) were injected into the hemocoel with GN6 or chitin at doses of 2.5, 1.3, 0.63, 0.31, and 0.15 µg per larva. Four hours after injection, the larvae were challenged with *Pseudomonas aeruginosa* (PAO1 strain) and maintained at 37°C. Survival was assessed 18 hours post-infection, and dose– response curves were fitted using a nonlinear regression model to calculate the median effective dose (ED₅₀). GN6 and chitin were found to increase larval survival in a dose-dependent manner, with calculated ED₅₀ values of 0.62 µg/larva and 0.48 µg/larva, respectively.

To investigate the structure–activity relationship, chitin oligosaccharides with lower degrees of polymerization (*N*-acetylglucosamine (GlcNAc, GN1), chitobiose (GN2), chitotriose (GN3), chitotetraose (GN4), and chitopentaose (GN5)) were injected under the same conditions. These oligosaccharides were purchased from Tokyo Chemical Industry Co., Ltd. (TCI, Tokyo, Japan). The infection resistance conferred by each compound was evaluated by comparing post-infection survival rates with those of GN6 and chitin.

### Statistical Analysis

All statistical analyses (except RNA-seq analysis) were performed using RStudio (version 2024.12.0+467; R Foundation for Statistical Computing, Vienna, Austria) with the following R packages: *survival*, *survminer*, *ggplot2*, *dplyr*, and *broom*. Kaplan–Meier survival curves were generated and compared using the log-rank test. Dose–response curves were analyzed by nonlinear regression to calculate the median effective dose (ED₅₀). For luciferase reporter assays, one-way ANOVA followed by Tukey’s multiple comparison test was used. A p-value < 0.05 was considered statistically significant.

## ACKNOWLEDGEMENTS

This work was supported in part by JSPS KAKENHI Grant Number 20K16253 and 22K15461 (to AM); the Open Innovation Research and Application Promotion Program (JPJ011937, Research Project No. 05001a1a2) from the Bio-oriented Technology Research Advancement Institution (BRAIN) (to AM); the Morinomiyako Medical Research Foundation (to AM); the Takeda Science Foundation (to AM); the Uehara Memorial Foundation (to AM); a research grant from Teikyo University (to AM); and the Life Science Foundation of Japan (to AM).

## Supplemental Table & Figure

**Supplementary Figure S1.**
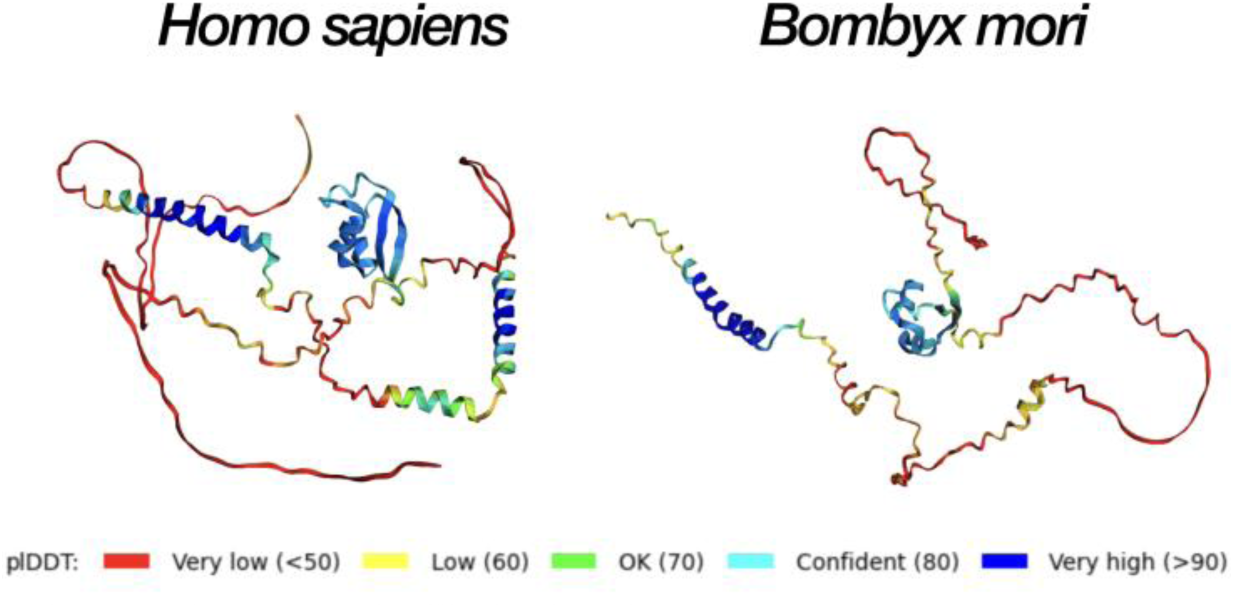
AlphaFold-predicted structures and confidence scores for LysM-containing proteins from *Homo sapiens* and *Bombyx mori*. Predicted structures of human LYSMD3 and its putative *B. mori* homolog (XP_004933441.1) were generated using AlphaFold and visualized with per-residue confidence scores (pLDDT). Regions with high prediction confidence (pLDDT > 90, shown in dark blue) were concentrated within the LysM domain in both species, supporting the structural comparison presented in Figure 1B. Regions outside the LysM domain showed lower confidence scores (light blue to red) and were excluded from RMSD-based comparative analysis.

Supplemental Table 1

The annotated list of differentially expressed genes (DEGs) is provided in Supplementary Table 1 (attached as a separate file due to file size limitations).

**Supplementary Figure S2.**
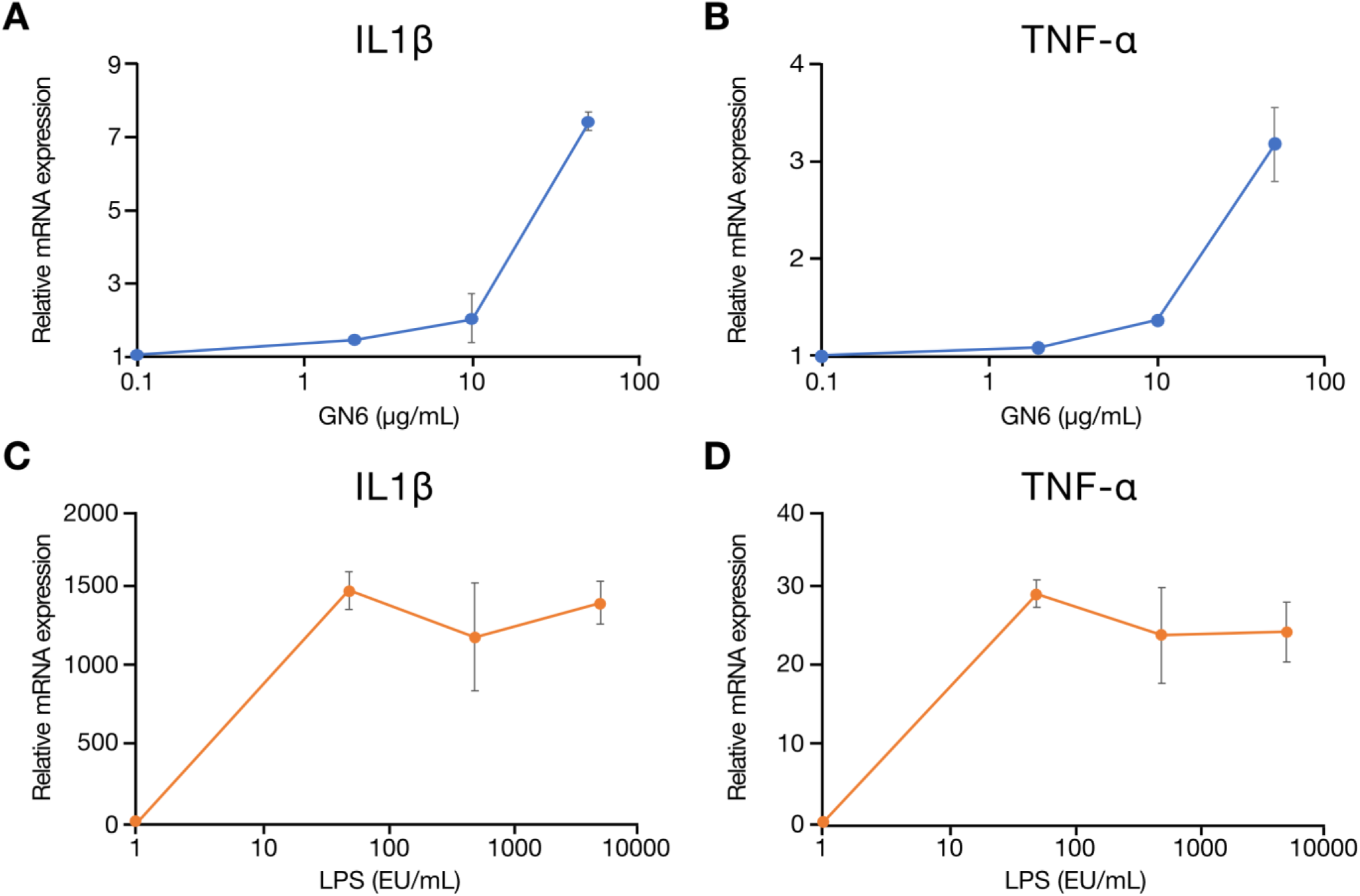
GN6 induces the expression of proinflammatory cytokine genes in RAW264.7 cells. Mouse macrophage RAW264.7 cells were stimulated with GN6 (A, B) or lipopolysaccharide (LPS, C, D) for 2 h. Gene expression levels of the proinflammatory cytokines *Il1b* and *Tnf* were analyzed by qRT-PCR and normalized to the internal control gene *Rps18*. Results are presented as fold changes relative to unstimulated controls. GN6 induced dose-dependent upregulation of both cytokine genes, whereas LPS induced a strong response in both *Il1b* and *Tnf*. Data represent mean ± SD (n = 3 each).

**Supplementary Figure S3.**
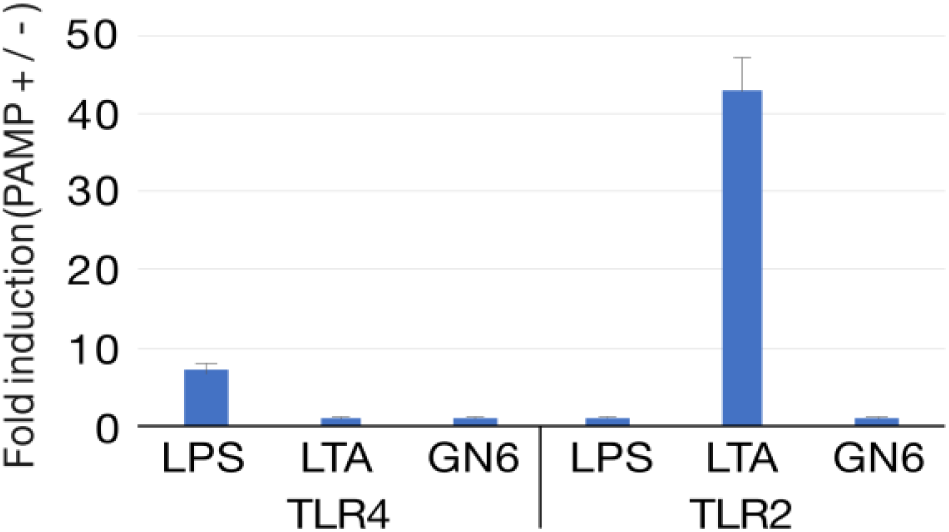
Evaluation of GN6-induced activation of human TLR2 and TLR4 pathways using HEK293 reporter cells. HEK293 reporter cells expressing either human TLR4/CD14/MD2 or TLR2, along with a firefly luciferase gene under the control of an NF-κB response element, were used to assess GN6-induced pathway activation. Cells were stimulated for 6 h with GN6 (5 µg/mL), lipopolysaccharide (LPS; 1 µg/mL) as a positive control for TLR4, or lipoteichoic acid (LTA; 1 µg/mL) as a positive control for TLR2. Luciferase activity was quantified using the Dual-Luciferase Reporter Assay System, and firefly luciferase signals were normalized to Renilla luciferase. Results are expressed as fold induction relative to unstimulated controls. Data represent the mean ± SD of three replicates.

**Supplementary Figure S4.**
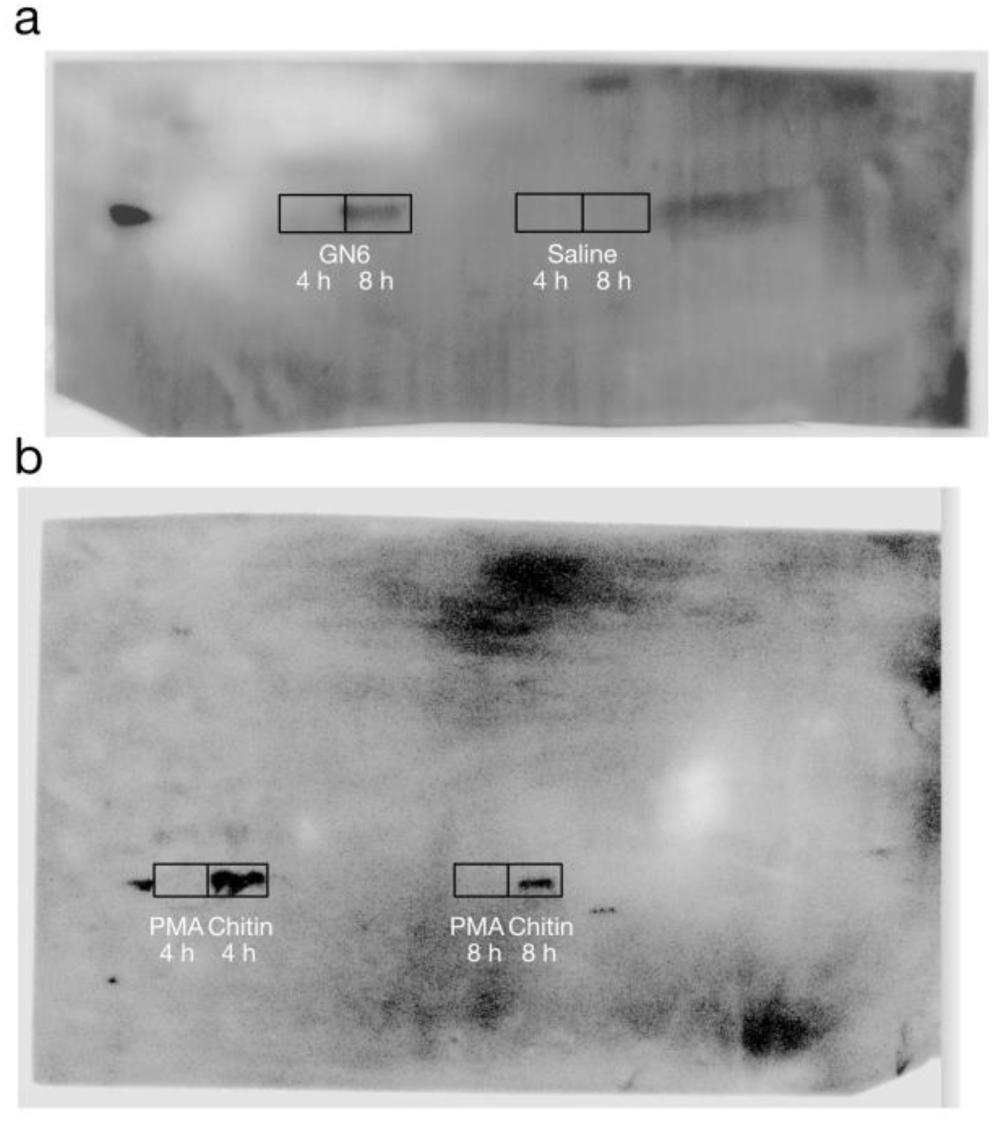
Full Western blot images for detection of Cecropin B in silkworm hemolymph following treatment with GN6, chitin, or PMA. (a, b) Fifth-instar day-2 *Bombyx mori* larvae were injected with GN6 (2.5 μg/larva), chitin (2.5 μg/larva), PMA (0.5 ng/larva), or saline, and hemolymph samples were collected at 4 h and 8 h post-injection at 27°C. Cecropin B protein levels in the hemolymph were analyzed by western blotting. (a) Blot showing samples from larvae treated with GN6 and saline. (b) Blot showing samples from larvae treated with PMA and chitin. The boxed areas indicate regions cropped and presented in the main text (Figure 3C).

